# Individual differences in post-encoding sleep continuity predict context memory accuracy and supporting ERPs in younger and older adults

**DOI:** 10.64898/2026.07.06.736892

**Authors:** Chuu Nyan, Aiden Wachnin, Soroush Mirjalili, Sahana Ram, Masoud Seraji, Audrey Duarte

**Author notes:** **Corresponding Author:** Chuu Nyan, B.S., Research Associate, The University of Texas at Austin, Department of Psychology.

## Abstract

Post-encoding sleep plays an essential role in episodic memory consolidation. Much of the existing literature on sleep and memory relies on deprivation paradigms or laboratory-controlled sleep. Relatively few studies have examined how naturalistic post-encoding sleep relates to memory retrieval and its supporting neural activity, or whether age-related impairments in this sleep are linked to those in episodic memory. In the present study, we used actigraphy and electroencephalography to examine how post-encoding sleep quality relates to context memory performance and retrieval-related ERPs supporting performance in younger and older adults. Participants encoded object–scene pairs and were tested on matching and mismatching pairs after a 96-hour sleep-filled delay. We found that greater post-encoding sleep continuity predicted better delayed context memory performance for mismatching pairs across age groups. Post-encoding sleep continuity was also associated with larger ERP differences between context hits and misses for context-matching pairs, for ERP effects associated with post-retrieval monitoring operations across age groups. Together, these findings suggest that more continuous, naturalistic post-encoding sleep facilitates episodic memory performance and neural mechanisms supporting episodic memory retrieval across adult age.

## Introduction

Sleep is essential for memory consolidation (for reviews see, Diekelmann & Born, 2010; Rasch & Born, 2013). Previous studies using sleep deprivation techniques have demonstrated that sleep disruption in the period post-encoding can impair episodic memory performance, particularly in younger adults (for a review see, Newbury et al., 2021). Evidence in older adults, who tend to have worse habitual sleep (e.g., shorter total sleep time (TST) and decreased sleep efficiency) and reduced episodic memory performance, is mixed (Lo et al., 2016; Mander et al., 2017; Sümer & Kaynak, 2025). Understanding the relationship between sleep and memory in older age is important, particularly given that poor habitual sleep is a risk factor for Alzheimer’s Disease (for a review see, Han et al., 2024).

Much of the existing literature has manipulated sleep in controlled laboratory environments, which fails to capture individual differences in naturalistic sleep (Morrow et al., 2023). Actigraphy provides multi-night estimates of various sleep parameters such as TST, and sleep quality measures like sleep efficiency, and wakefulness after sleep onset (WASO). Previous work has demonstrated that greater average and night-to-night variability in sleep quality is linked to worse episodic memory performance across age (for a review see, Hokett et al., 2021), and reduced retrieval-related neural activity (Hokett et al., 2022; Hokett & Duarte, 2019). It is unknown, however, whether individual differences in sleep quality during the post-encoding/consolidation period, per se, influence episodic memory performance and neural mechanisms supporting successful retrieval. While prior research links individual differences in naturalistic sleep quality and episodic memory performance across the adult lifespan (for a review see, Hokett et al., 2021), the absence of a defined sleep-filled post-encoding period has precluded addressing this question.

In the present study, we investigated the link between post-encoding sleep quality and episodic memory and supporting retrieval neural mechanisms in younger and older adults. We examined recognition for items and item-context pairs, presented as matching (intact) or mismatching (recombined), compared to their presentation during encoding. While matching item-context pairs may be recognized on the basis of familiarity, memory for mismatching pairs involves recollecting the relationship between items and their previously associated contexts (Yonelinas, 2002; Curran, 2000). The neural mechanisms supporting familiarity and recollection have been extensively examined using event-related potentials (ERPs) (Duarte et al., 2006; Harand et al., 2012; Wang et al., 2012). The FN400 (~300-500 ms post-stimulus) is linked to familiarity-based recognition, differentiating between items that people recognize, regardless of context memory accuracy, and those they correctly reject as new (for reviews see, Curran, 2000; Friedman & Johnson, 2000; Rugg & Curran, 2007), and is typically preserved with age (Koen & Yonelinas, 2016). A posterior old-new effect (~400-800 ms post-stimulus) is typically observed as greater positivity over posterior electrodes for correctly recognized items, especially when accompanied by successfully retrieved episodic details (for reviews, Friedman & Johnson, 2000; Rugg & Curran, 2007). The late frontal old-new effect (onsetting ~1000 ms post-stimulus and sustained until response) is common in context memory studies in which post-retrieval monitoring of specific, objective features is needed to make a decision (Cruse and Wilding, 2009; Friedman and Johnson, 2000; Senkfor and Van Petten, 1998; Wilding and Rugg, 1996). The late posterior negativity (LPN), observed in a similar late time period as the late frontal old-new ERP, is associated with memory reconstruction, particularly evident in tasks requiring retrieval of perceptual contextual features (for a review see: Mecklinger et al., 2016).

We recruited a local community sample of younger and older adults to examine the contribution of individual differences in habitual sleep patterns, particularly during post-encoding sleep, to episodic memory retrieval after a 96-hour delay and supporting neural activity. We assessed the extent to which naturalistic, post-encoding sleep quality and quantity relate to neural activity supporting context memory retrieval after the delay in younger and older adults. Given evidence showing that memories depending on recollection may be more sensitive to sleep than those supported by familiarity alone (Newbury et al., 2021; Hokett et al., 2022), we predicted that greater post-encoding sleep quality would be associated with better context memory accuracy across age, particularly for mismatching pairs, and for ERPs linked to recollection and/or post-retrieval monitoring and reconstruction, which are often reduced in older age (Koen & Yonelinas, 2014; Kwon et al., 2023).

## Methods

### Participants

Eighty-five participants were recruited from the greater Austin area through mailings, phone calls, and flyer distributions. Nine participants were excluded due to missing sleep data resulting from the accelerometer malfunction. One additional participant was excluded from subsequent analyses due to insufficient sleep data (less than one night), due to forgetting to wear the accelerometer during the post-encoding sleep period. As a result, seventy-five participants were included in principal component analyses (PCA) of the sleep data. An additional five participants were excluded from the analysis: one due to having excessive sweat-related electroencephalography (EEG) artifact, two for experimenter task administration error, and two for providing insufficient correct responses during retrieval (fewer than 10 correct trials). The final sample consisted of 70 participants, including 35 younger adults (22 females, 12 males, 1 self-reported as other gender; age range: 18–30 years) and 35 older adults (16 females, 19 males; age range: 56–79 years). Our primary effects of interest were associations between sleep quality and memory performance, and between sleep quality and EEG measures of retrieval success. Based on our prior study (Hokett E and Duarte A, 2019) showing medium effect sizes for regression models including younger and older adult samples with habitual sleep quality predicting associative memory performance (R2 = 0.09) and EEG differences between successfully vs. unsuccessfully remembered pairs (R2 = 0.12), we estimated a total sample of 67 participants for the moderated regression models. Thus, our sample of 70 is sufficiently powered. None of the participants reported neurological disorders, psychiatric conditions, sleep disorders, or cardiovascular disease. Younger adults self-reported race/ethnicity: 22.9% identified as Non-Hispanic White, 40% as Asian, 2.9% as Black, 20% as Hispanic, 8.57% as multiracial, and 5.71% did not report. Older adults self-reported race/ethnicity: 68.6% identified as Non-Hispanic White, 2.86% as Asian, 5.71% as Black, 20% as Hispanic, and 2.86% did not report. Participants also self-reported years of education, right-handedness, and vision (normal or corrected). All participants signed consent forms approved by the Institutional Review Board of the University of Texas at Austin. Participant demographics and neuropsychological test scores are reported in Table 1.

**Table 1.**
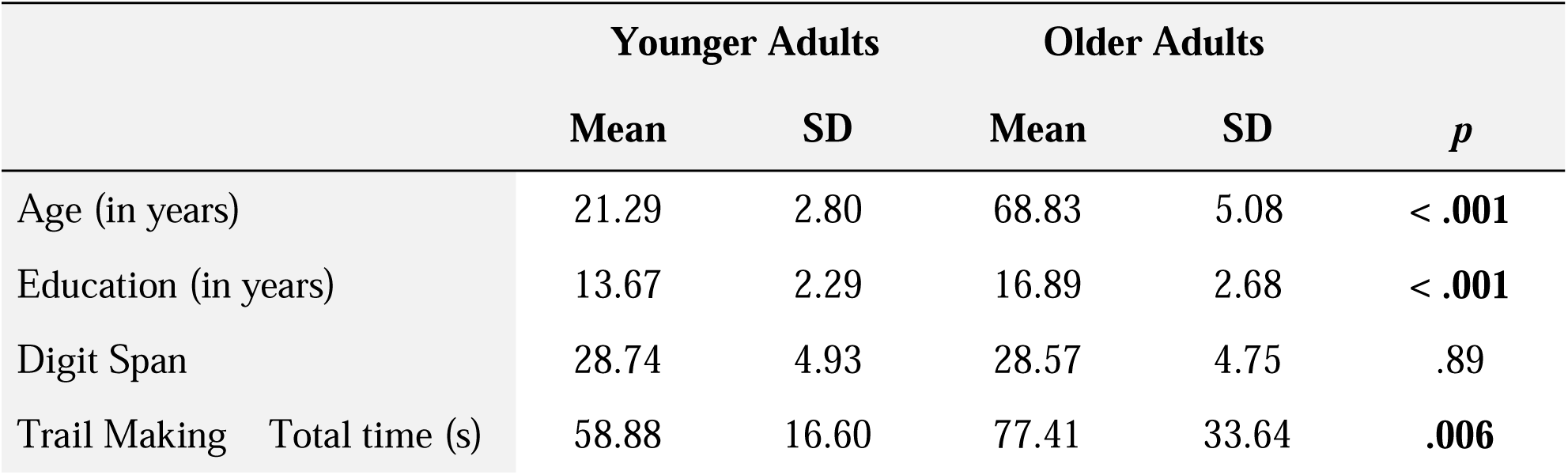
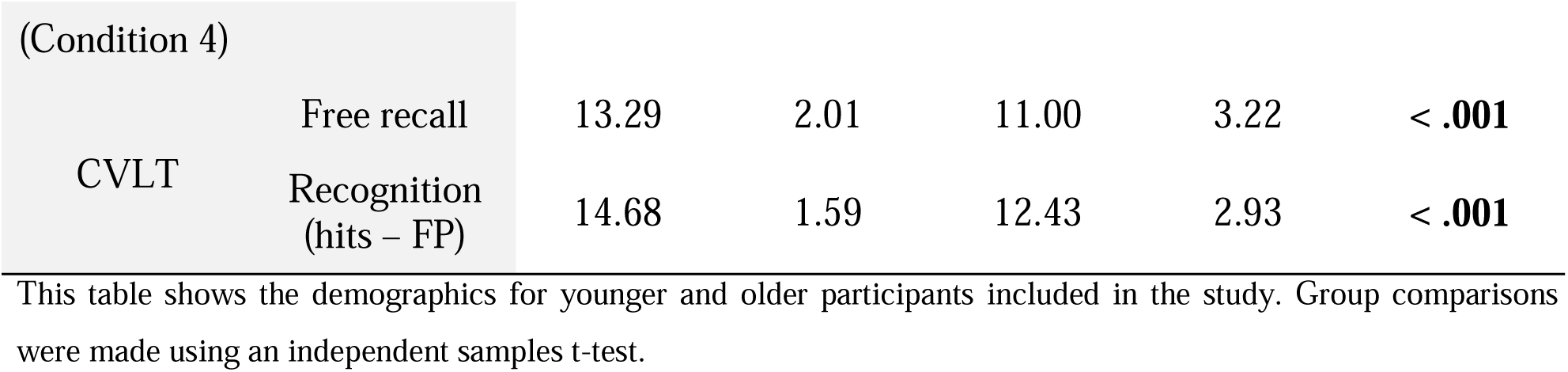
Participant demographics and neuropsychological tests raw scores for the younger and older age groups.

|  | Younger Adults |  | Older Adults |  | *p* |
| --- | --- | --- | --- | --- | --- |
|  | Mean | SD | Mean | SD |  |
| Age (in years) | 21.29 | 2.80 | 68.83 | 5.08 | < .001 |
| Education (in years) | 13.67 | 2.29 | 16.89 | 2.68 | < .001 |
| Digit Span | 28.74 | 4.93 | 28.57 | 4.75 | .89 |
| Trail Making Total time (s) | 58.88 | 16.60 | 77.41 | 33.64 | .006 |

|  |  |  |  |  |  |  |
| --- | --- | --- | --- | --- | --- | --- |
| (Condition 4) |  |  |  |  |  |  |
| CVLT | Free recall | 13.29 | 2.01 | 11.00 | 3.22 | < .001 |
|  | Recognition (hits – FP) | 14.68 | 1.59 | 12.43 | 2.93 | < .001 |

This table shows the demographics for younger and older participants included in the study. Group comparisons were made using an independent samples t-test.

### Study materials

We obtained 288 object images and 12 scene images from Google Images and the Bank of Standardized Stimuli (BOSS). Each trial consisted of an object–scene pair, with a namable object superimposed on a namable scene presented on a neutral gray background, as seen in Figure 1. Scene images measured 15 × 15 cm, and object images measured 4 × 4 cm. Both images were centered on the screen to ensure presentation within the central visual field. At a viewing distance of 70 cm, scene images subtended a visual angle of approximately 12.2°, and object images subtended a visual angle of approximately 3.2°. Visual angles were held constant across trials.

**Fig. 1.**
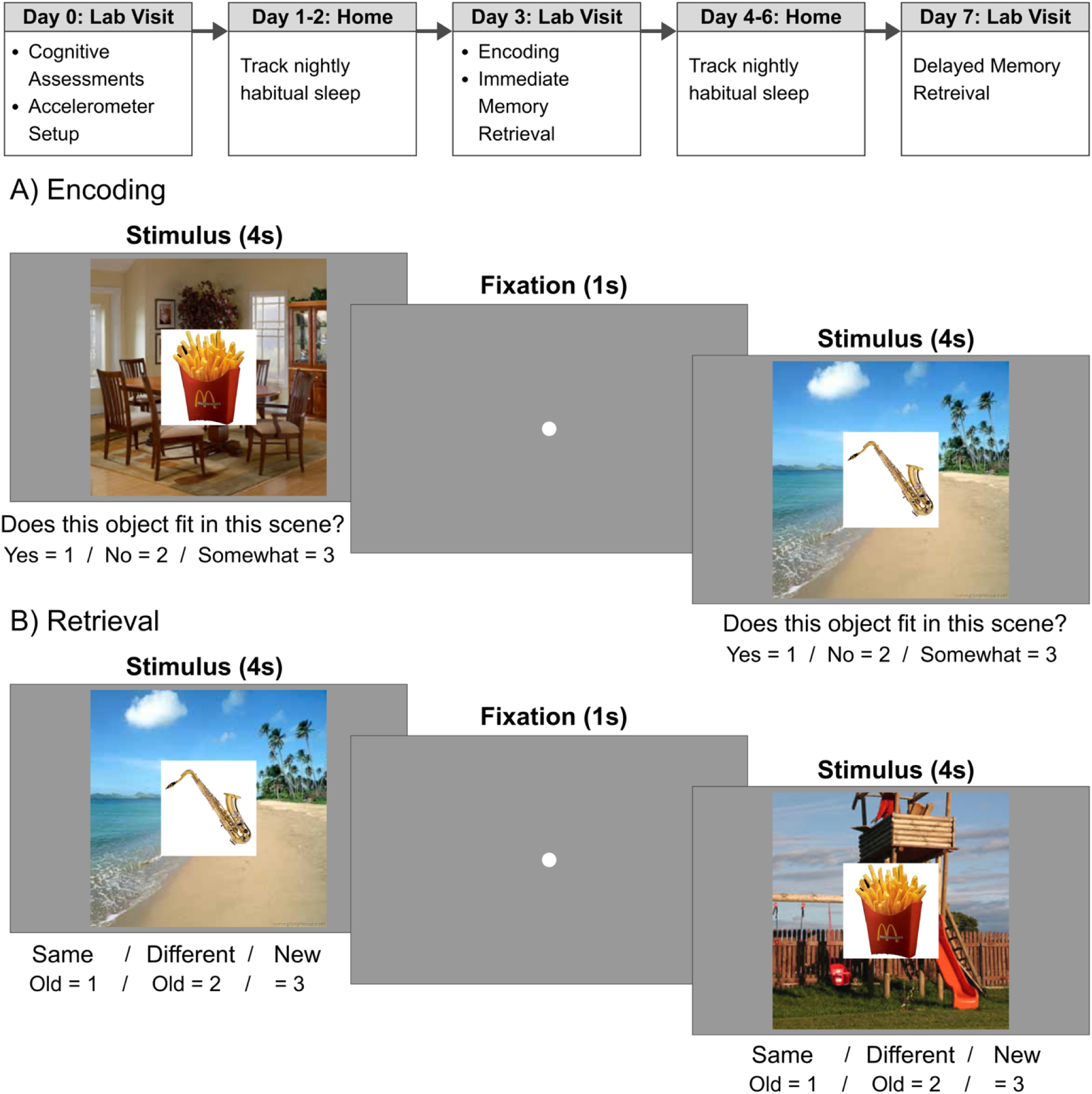
Experimental Procedure.

### Study procedure

The 7-day procedure can be seen in Figure 1. During the first lab visit, participants completed a battery of neuropsychological assessments to screen for mild cognitive impairment. Participants were also provided with an accelerometer (Philips Respironics Actiwatch 2) and a sleep diary to monitor participants’ bedtime and wake time to validate actigraphy-derived nightly sleep information. They were asked to wear the Actiwatch2 at all times, except during bathing or swimming. The neuropsychological battery included the California Verbal Learning Test–Third Edition (CVLT-3; Delis et al., 2017), the Delis–Kaplan Executive Function System (D-KEFS), Trail Making Test (Delis et al., 2001), and subtests from the Wechsler Adult Intelligence Scale–Fourth Edition (WAIS-IV; Wechsler, 2008), which assessed working memory and executive function. Participants returned to the lab on Day 3 and Day 7 of the study to complete a context memory task (see Figure 1 for an overview of the task procedures). All participants completed memory encoding and immediate retrieval on Day 3, followed by delayed retrieval on Day 7. The participants were not informed that a follow-up memory test would be conducted. EEG was recorded during all task phases.

Context memory task stimuli, screen prompts and timings are shown in Figure 1. During encoding, 288 object–scene pairs were presented, and participants were instructed to evaluate whether the object at the center of the display fit within the context of the background scene (i.e., “Does this object fit in this scene?”), and to press one of 3 options using a keyboard: (1) yes, (2) no, (3) somewhat. Half of the pairs were designed to have matching object-context associations (e.g. stapler in an office) and the other half non-matching object-context associations (e.g. saxophone on a beach). Object-scene pairs were presented in six blocks of 72 trials each per encoding block. Immediately following the encoding, participants completed an immediate retrieval task. For each retrieval tasks, 432 object-scene pairs were presented. Among them, 144 matching pairs were presented in which the studied object was presented with the same scene background as during encoding, along with 144 mismatching pairs in which the studied object was presented with a different scene background, and 144 new pairs in which new objects were presented with one of the previously studied scenes. These pair types were presented in a randomized order in six blocks of 72 trials, with 24 trials per pair type per retrieval block. Participants indicated whether each object had been previously seen, and, if so, whether it had been paired with the same or a different background. Responses were made using one of three keys: (1) old object paired with the same background, (2) old object paired with a different background, or (3) new object. Participants returned to the lab after 96 hours to complete a delayed retrieval task. Immediate and delayed retrieval tasks consist of unique new object pairs from each other.

### EEG data acquisition and preprocessing

The EEG data was recorded using a 32-channel BrainProducts actiChamp system, operating at a 500 Hz with a 24-bit resolution. Electrode placement followed the standard 10-20 system, with Cz channel serving as the online reference and Fpz as the ground electrode (Jasper, 1958). Electrode positions included: C3, C4, CP1, CP2, CP5, CP6, Cz, F3, F4, F7, F8, FC1, FC2, FC5, FC6, FP1, FP2, FT9, FT10, Fz, O1, O2, Oz, P3, P4, P7, P8, Pz, T7, T8, TP9, and TP10. Additionally, four electrodes were used to monitor eye movements: two for horizontal electrooculogram (HEOG) at the outer canthi of both eyes, and two for vertical electrooculogram (VEOG) above and below the right eye.

EEG analyses were focused on delayed retrieval to test our hypotheses of interest. All offline data analysis was performed using the EEGLAB toolbox v2023.0 (Delorme A & Makeig S, 2004) under MATLAB R2023b. EEG data was re-referenced to the average of the left (TP9) and right (TP10) mastoid electrodes and downsampled to 250 Hz. We then applied a band-pass filter between 0.05 Hz and 80 Hz frequencies to the continuous data. Epochs were extracted between 300 ms before and 2000 ms after the onset of the stimulus, and baseline corrected using the 300 ms before the stimulus onset. We visually inspected and rejected epochs with non-ocular artifacts (e.g., muscle, electrode, and sweat artifacts). In the case of noisy electrodes (e.g, at least 40% of the epochs are rejected due to one noisy electrode), we interpolated using the surrounding electrodes to estimate the activity of the electrode that is being interpolated. After this inspection, we applied Independent Component Analysis (ICA) to identify components that were associated with eye movement (e.g., eye blinks or horizontal eye movements), muscle, or heart artifacts with >0.9 probability. Following ICA, epochs with voltages exceeding ±150 microvolts were removed. Lastly, to reduce residual high frequency noise, we applied a lowpass filter of 20 Hz to all individual averages before averaging across all participants.

### Sleep measurements and principal component analysis

From the actigraphy recordings, we obtained total sleep time (TST–minutes asleep), sleep efficiency (percentage of time in bed spent asleep), wake after sleep onset (WASO–total number of minutes spent awake after initial sleep onset), sleep fragmentation index (the sum of sleep bouts under 60s with physical movement and percent physical movement during sleep) and number of awakenings (the total number of wake bouts during the sleep period). Our main hypotheses were related to the post-encoding sleep period (4 nights). For these nights, we computed both the mean of each sleep measure. We additionally computed means for the pre-encoding (3 nights) to conduct exploratory analyses (see Supplementary Materials).

PCA with Varimax rotation was conducted to reduce the dimensionality of the sleep measures and to extract latent sleep components. Components with eigenvalues greater than 1 were retained. Two components were identified for each model: a sleep time component, characterized by TST, and a sleep continuity component, characterized by sleep fragmentation index, sleep efficiency, number of awakenings, and WASO (see Table 2 and Supplemental Table 1).

**Table 2.** This table shows component loadings for the post-encoding sleep variables.

| Mean Sleep Variables | Post-Encoding Sleep (96 hrs) |  |
| --- | --- | --- |
|  | Sleep Continuity | Sleep Time |
| Total Sleep Time | 0.11 | **0.96** |
| Sleep Efficiency | **0.84** | 0.22 |
| Number of Awakenings | **-0.71** | 0.59 |
| Wake After Sleep Onset (WASO) | **-0.80** | 0.50 |
| Sleep Fragmentation Index | **-0.87** | -0.001 |

Bolded text indicates the strongest variables for each principal component. Sleep continuity explained 62% variance; sleep time explained 38% of the sample variance. Model fit ( $\chi^2(1) = 32.360, df = 1, p < .001$ ).

### Sleep and memory performance analysis

Context memory performance was measured by first calculating the response proportions for each response category: context hits were calculated as the proportion of matching pairs endorsed as old/same, context misses were calculated as the proportion of matching pairs endorsed as old/different, context correct rejection were calculated as the proportion of mismatching pairs endorsed as old/different, and context false alarms were calculated as the proportion of mismatching pairs endorsed as old/same. Memory retention was calculated as the difference between immediate and delayed context memory retrieval, with more negative values indicating greater memory decline over time.

We used hierarchical multiple regressions to test the hypotheses that post-encoding sleep quality was associated with delayed matching and mismatching context memory performance, potentially to a greater degree for older adults. To this end, we entered age and post-encoding sleep PCA components as predictors in Block 1, followed by the interaction between age and these components in Block 2. Post-encoding sleep continuity and post-encoding sleep time were used as predictor variables in separate models. We explored the same models for retention memory using pre-encoding and post-encoding sleep as predictors, as shown in Supplementary Tables 2 and 3.

### ERP old-new effect analysis

The current study analyzed the EEG data from the delayed retrieval phase. We averaged stimulus-locked ERP waveforms separately for context hits, context misses, context correct rejections, context false alarms, and correctly rejected new objects (item correct rejections). Old object-scene pairs for which subjects responded “New,” indicating that the old object was forgotten (item misses) and falsely recognized new object trials (item false alarms) were not of interest for our hypotheses and were also not included in ERP analyses.

We used a mass-univariate permutation test to examine ERP old-new effects in younger and older adults. This approach corrects for multiple comparisons across ROIs and time points, controls the family-wise error rate, and yields robust significance estimates. Instead of evaluating effects from individual data points, this method performs a cluster-based permutation test across multiple channels and time points. The analysis groups adjacent data points in space and time and estimates the probability of observing the clusters by chance. The ERP data were downsampled to 125Hz for this analysis.

To investigate the reliability of old-new effects in each age group, we performed a 2 x 2 between-subject analysis of variance (ANOVA) with factors of Condition x Age (younger, and older). Separate models were conducted for matching context pairs (Context Hit, Context Miss, Item Correct Rejection) and mismatching context pairs (Context Correct Rejection, Context False Alarm, Item Correct Rejection). Follow up pairwise ANOVAs were conducted for significant clusters to identify the source of the main effects and/or interactions.

### Sleep and context memory ERP analysis

We used hierarchical multiple regressions to investigate whether individual differences in sleep quality, age, and their interaction predicted differences in voltages between context memory conditions at the delayed retrieval. We entered age (Younger and Older) and sleep variables as predictors in block 1, and the interaction between age and sleep variables in block 2. Voltage differences between matching (context hits vs. context misses) and mismatching (context correct rejections vs. context false alarms) memory conditions at delay retrieval were used as predictor variables in the separate models. In order to reduce the number of comparisons while focusing on scalp regions where the ERPs of interest are typically observed, we divided electrodes into 4 regions of interest (ROIs): Left Frontal (F3, FC1, FC5), Right Frontal (F4, FC2, FC6), Left Posterior (CP1, CP5, P3), and Right Posterior (CP2, CP6, P4). Using a sliding time window approach, we divided the 2000-ms retrieval period into nineteen 200-ms time windows where the consecutive time windows were 100-ms apart from each other (i.e., [0-200 ms], [100-300 ms], [200,400 ms], …, [1800-2000 ms]). We next found the time windows where the voltage amplitude contrast was significantly predicted by the sleep variables, age, and the interaction between sleep and age. If adjacent 200-ms time windows showed significant effects, they would be *clustered* together into larger time-windows (e.g., if [300-500 ms], [400-600 ms], [500-700 ms] showed significant effects, we would create a significant cluster of [300-700 ms]). We next performed a multiple comparison permutation test to obtain robust significance estimates and control the false discovery rate of the significant correlations. To elaborate, for each 1000 permutations, we shuffled the voltage contrast values over different 200-ms windows (e.g., changing the ERP contrast from [700-900 ms] to the ERP contrast from [300-500 ms]) and identified the clusters that had a significant relationship with the sleep component. For each permutation, we measured the *duration* of the longest cluster with a significant effect (e.g., 200-ms, 300-ms, etc.). After repeating this process 1000 times, we obtained a null distribution of the cluster lengths. If any of our original clusters were longer than at least 95% of the cluster lengths from the null distribution, we would keep them for our subsequent analyses. Memory-related significant voltage clusters will be called matching or

## Results

### Principal components analysis for sleep variables

The principal component analyses yielded two sleep components. Table 2 shows principal component loadings for two post-encoding sleep components: Sleep Continuity (Sleep Fragmentation Index, Sleep Efficiency, Number of Awakening and WASO) and TST. Independent samples t-tests revealed that younger adults (Mean = 0.26, SD = 0.47) compared to older adults (Mean = −0.17, SD = 0.52) exhibited significantly better sleep continuity during the post-encoding sleep (t(68) = 3.61, p < .001). There were no significant group differences between sleep time during the post-encoding sleep (younger adults: Mean = −0.05, SD = 1.06; older adults: Mean = 0.11, SD = 0.95; t(68) = −0.66, p = .51). Supplemental Table 1 includes PCA results for 7-nights and pre-encoding sleep.

### Behavioral results

#### Greater context memory performance for context matching than mismatching pairs and for younger than older adults

Table 3 presents the mean proportion of item hits and misses, context hits and misses, and context correct rejections and false alarms for each age group. Context memory accuracy, calculated as the proportion of p(context hits)/[p(context hits)+p(context misses)] for matching pairs and p(context correct rejection)/[p(context correct rejection)+p(context false alarms] for mismatching pairs, is shown for the immediate and delayed retrieval tasks, along with retention scores, in Figure 2A. Memory retention was calculated as the difference between delayed and immediate memory retrieval. Younger and older adults performed above chance (0.5) on context memory retrieval tasks for both immediately and after a delay.

**Fig. 2.**
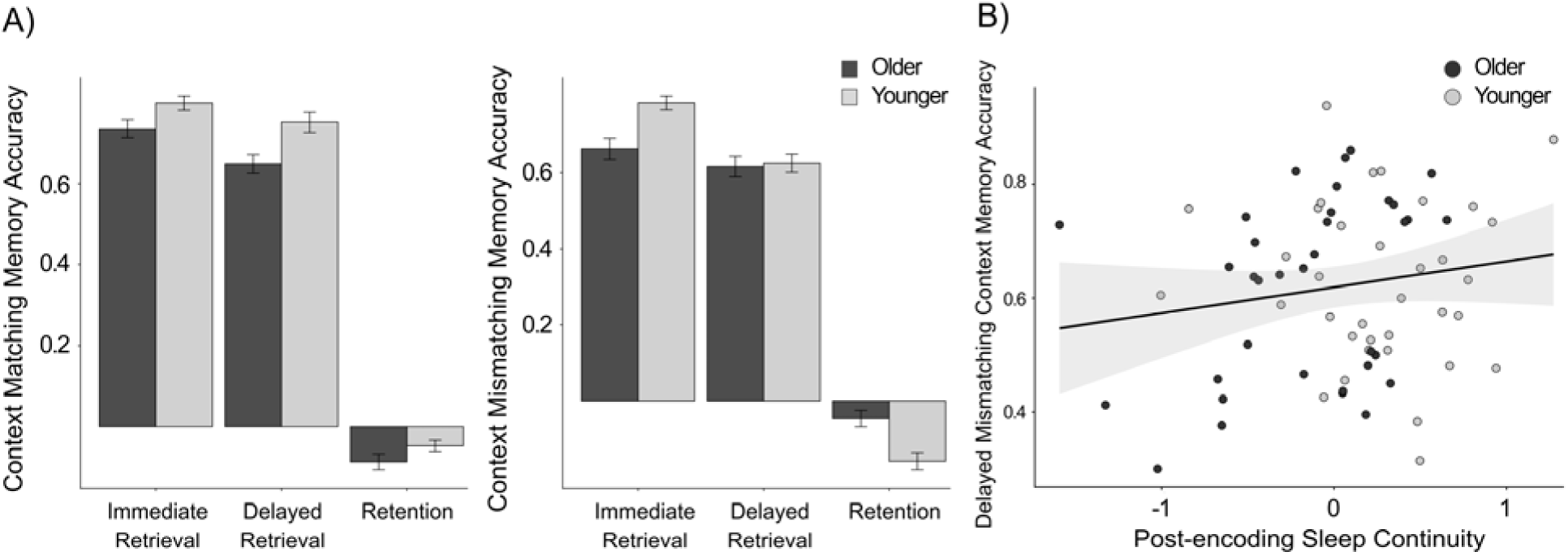
A) Proportions of context matching and mismatching accuracy for younger and older adults. Error bars represent standard error of the mean. B) Scatter plot showing the significant relationship between the post-encoding sleep continuity PCA component score and delayed context mismatching memory accuracy for younger and older adults.

**Table 3.** Mean proportions of memory response categories by study history and pair type for younger and older adults.

| **Context matching pairs** |  |  |  |  |  |  |
| --- | --- | --- | --- | --- | --- | --- |
|  | Memory response categories | Younger Adult | Older Adult | *t* | *p* | *Cohen's d* |
| Immediate | Item Hit | 0.84 (0.10) | 0.78 (0.13) | 2.06 | .04 | 0.49 |
|  | Item Miss | 0.14 (0.10) | 0.18 (0.11) | -1.67 | .10 | 0.40 |
|  | Context Hit (of Item Hits) | 0.80 (0.10) | 0.73 (0.13) | 2.27 | .03 | 0.54 |
|  | Context Miss (of Item Hits) | 0.20 (0.10) | 0.27 (0.13) | -2.27 | .03 | 0.54 |
| Delayed | Item Hit | 0.78 (0.12) | 0.72 (0.12) | 1.04 | .05 | 0.47 |
|  | Item Miss | 0.16 (0.09) | 0.18 (0.12) | -1.04 | .28 | 0.26 |
|  | Context Hit (of Item Hits) | 0.75 (0.15) | 0.65 (0.14) | 2.69 | .004 | 0.72 |
|  | Context Miss (of Item Hits) | 0.25 (0.15) | 0.35 (0.14) | -2.77 | .004 | 0.72 |
| **Context mismatching pairs** |  |  |  |  |  |  |
| Immediate | Item Hit | 0.75 (0.14) | 0.65 (0.15) | 2.80 | .01 | 0.67 |
|  | Item Miss | 0.20 (0.11) | 0.26 (0.12) | -2.47 | .02 | 0.59 |
|  | Context Hit (of Item Hits) | 0.78 (0.11) | 0.66 (0.17) | 3.62 | .001 | 0.87 |
|  | Context Miss (of Item Hits) | 0.22 (0.11) | 0.34 (0.17) | -3.62 | .001 | 0.87 |
| Delayed | Item Hit | 0.63 (0.15) | 0.58 (0.17) | 0.36 | .23 | 0.29 |
|  | Item Miss | 0.26 (0.14) | 0.27 (0.16) | -0.36 | .72 | 0.09 |
|  | Context Correct Rejection (of Item Hits) | 0.63 (0.14) | 0.62 (0.16) | 0.60 | .81 | 0.06 |
|  | Context False Alarm (of Item Hits) | 0.37 (0.14) | 0.38 (0.16) | -0.21 | .81 | 0.06 |
| **New pairs** |  |  |  |  |  |  |
| Immediate | Item Correct Rejection | 0.92 (0.09) | 0.88 (0.08) | 2.07 | .04 | 0.50 |
|  | Item False Alarm “Matching” | 0.02 (0.02) | 0.04 (0.04) | -1.94 | .06 | 0.46 |
|  | Item False Alarm “Mismatching” | 0.06 (0.07) | 0.09 (0.06) | -1.73 | .09 | 0.41 |
| Delayed | Item Correct Rejection | 0.82 (0.16) | 0.76 (0.16) | 1.62 | .11 | 0.39 |
|  | Item False Alarm “Matching” | 0.07 (0.08) | 0.10 (0.08) | -1.52 | .13 | 0.36 |
|  | Item False Alarm “Mismatching” | 0.11 (0.10) | 0.14 (0.12) | -1.25 | .22 | 0.30 |

This table shows the memory proportions for younger and older participants. Group comparisons were made using an independent samples t-test.

We conducted two-way ANOVAs for each memory delay including factors of Age (Younger, Older) and Pair Type (Matching, Mismatching). The ANOVA for immediate retrieval revealed main effects of Age [F(1, 136) = 17.92, p < .001, η^2^p = 0.12] and Pair Type [F(1, 136) = 3.87, p = .051, η^2^p = 0.030] but no interaction between these factors [F(1, 136) = 1.76, p = .19, η^2^p = 0.010]. For delayed retrieval, the ANOVA revealed main effects of Age [F(1, 136) = 5.01, p = .027, η^2^p = 0.040] and Pair Type [F(1, 136) = 9.90, p = .002, η^2^p = 0.070], but no significant interaction between these factors [F(1, 136) = 3.56, p = .061, η^2^p = 0.030]. As can be seen in Figure 3, context memory performance was greater for younger than older adults and for matching pairs than mismatching pairs across delays. The ANOVA for retention memory revealed main effects of Age [F(1, 136) = 3.39, p = .068, η^2^p = 0.020] and Pair Type [F(1, 136) = 3.25, p = .074, η^2^p = 0.020], and an interaction between these factors [F(1, 136) = 14.76, p < .001, η^2^p = 0.10]. The significant Age x Pair Type interaction in memory retention was driven by higher retention for mismatching pairs in older than younger adults [*t*(68) = −3.57, *p* = .001, Cohen’s *d* = 0.85] with no group difference in retention for matching pairs [*t*(68) = 1.66, *p* = .10, Cohen’s *d* = 0.40]. Retention was higher for match than mismatch pairs in younger [*t*(34) = 3.41, *p* = .002, Cohen’s *d* = 1.01] but not older adults [*t*(34) = −1.10, *p* = .28, Cohen’s *d* = 0.33].

**Fig 3.**
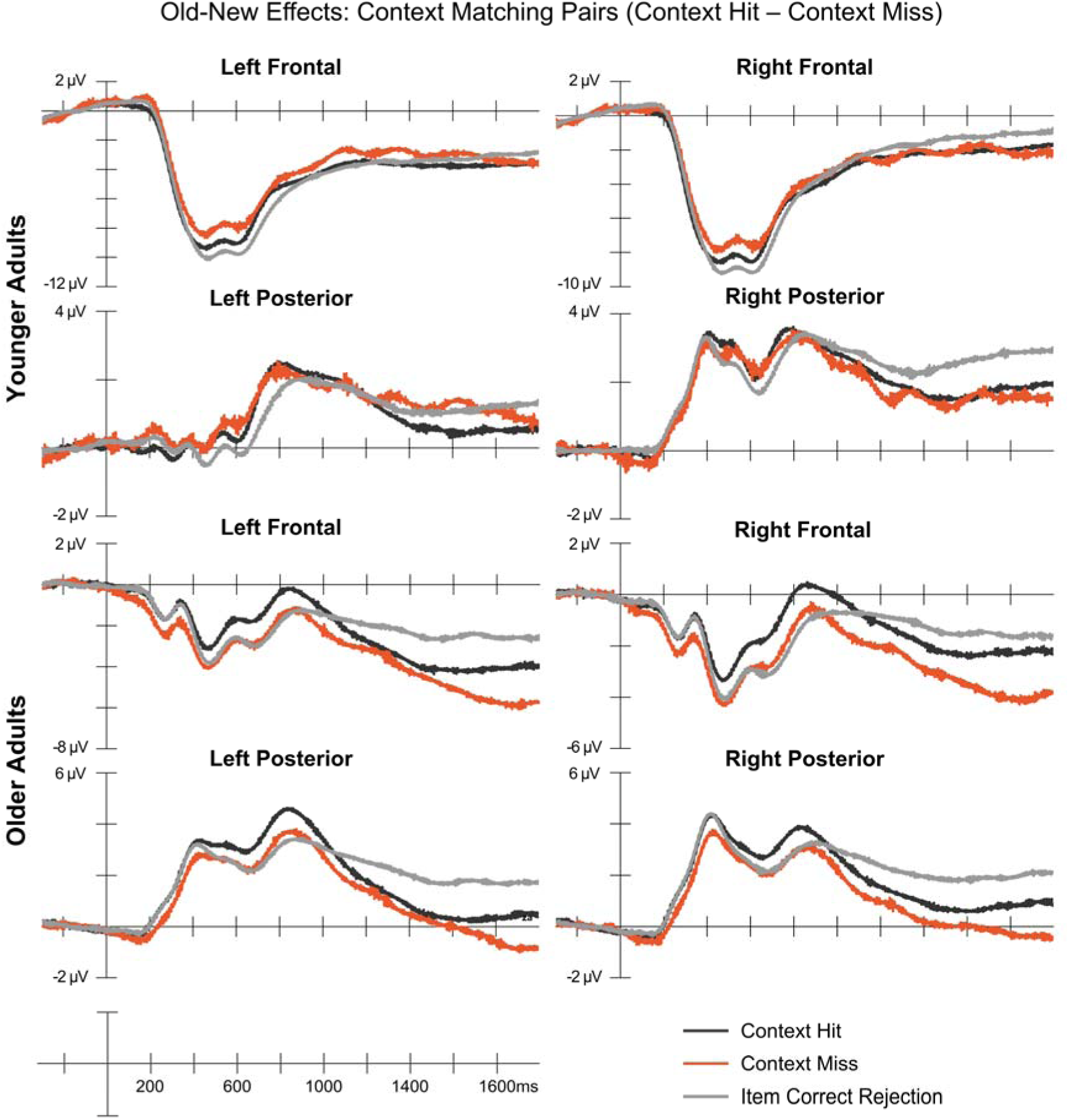
Grand ERPs for context matching pairs are depicted in this figure for each age group. Context matching pairs included an average number of 76.2 (range: 17–128) Context Hit, 32 (range: 6–90) Context Miss, 104 (range: 40–136) Item Correct Rejection.

#### Post-encoding sleep continuity is associated with context mismatching pair memory accuracy, across age

Results from hierarchical regression models examining relationships between sleep components and memory performance are shown in Table 4. A significant main effect of age was observed for context matching pairs, reflecting the reduced context memory accuracy for older than younger adults. There were no significant effects for the sleep components, however, for matching pair memory performance. By contrast, post-encoding sleep continuity was positively associated with context memory accuracy for mismatching pairs, across age groups (see Table 4 and Figure 2B). Sleep time was not significantly associated with either context memory accuracy measure. Therefore, sleep time was excluded from subsequent analyses.

**Table 4.** Summary of hierarchical multiple regression with age and post-encoding sleep predicting context matching and mismatching pair delayed context memory accuracy.

| **Outcome variable: delayed context matching memory** |  |  |  |  |  |  |
| --- | --- | --- | --- | --- | --- | --- |
| Predictor variables | **Block 1** |  |  | **Block 2** |  |  |
| | b | SE | $\beta$ | b | SE | $\beta$ |
| Intercept | 0.64 | 0.02 |  | 0.64 | 0.02 |  |
| Age | 0.12 | 0.04 | 0.42* | 0.12 | 0.04 | 0.41* |
| Sleep Continuity | -0.05 | 0.03 | -0.19 | -0.07 | 0.05 | -0.27 |
| Age x Sleep Continuity |  |  |  | 0.04 | 0.07 | 0.10 |
| $R^2$ | | 0.12 | | | 0.11 | |
| $\Delta R^2$ | | 0.12 | | | 0.005 | |
| F for $\Delta R^2$ | | 5.88 | | | 4.02 | |
| **Outcome variable: delayed context matching memory** |  |  |  |  |  |  |
| Intercept | 0.64 | 0.02 |  | 0.64 | 0.02 |  |
| Age | 0.11 | 0.03 | 0.35* | 0.11 | 0.03 | 0.37* |
| Sleep Time | 0.02 | 0.02 | 0.15 | 0.04 | 0.02 | 0.26 |
| Age x Sleep Time |  |  |  | -0.03 | 0.03 | -0.15 |
| $R^2$ | | 0.11 | | | 0.11 | |
| $\Delta R^2$ | | 0.11 | | | 0.01 | |
| F for $\Delta R^2$ | | 5.50 | | | 3.91 | |
| **Outcome variable: delayed context mismatching memory** |  |  |  |  |  |  |
| Intercept | 0.62 | 0.03 |  | 0.63 | 0.03 |  |
| Age | -0.01 | 0.04 | -0.04 | -0.004 | 0.04 | -0.02 |
| Sleep Continuity | 0.05 | 0.04 | 0.17 | 0.10 | 0.05 | 0.36* |
| Age x Sleep Continuity |  |  |  | -0.11 | 0.07 | -0.27 |
| $R^2$ | | -0.001 | | | 0.02 | |
| $\Delta R^2$ | | -0.001 | | | 0.03 | |
| F for $\Delta R^2$ | | 0.94 | | | 1.40 | |
| **Outcome variable: delayed context mismatching memory** |  |  |  |  |  |  |
| Intercept | 0.62 | 0.03 |  | 0.62 | 0.03 |  |
| Age | 0.01 | 0.04 | 0.03 | 0.01 | 0.04 | 0.03 |
| Sleep Time | 0.003 | 0.02 | 0.02 | 0.01 | 0.03 | 0.05 |
| Age x Sleep Time |  |  |  | -0.01 | 0.04 | -0.03 |
| $R^2$ | | -0.03 | | | -0.04 | |
| $\Delta R^2$ | | -0.03 | | | 0.0004 | |
| F for $\Delta R^2$ | | 0.05 | | | 0.04 | |

### EEG results

#### Mass univariate old-new ERP results

##### Matching context pairs

Mass univariate analysis results for matching context pairs are reported in Table 5. ERPs of context hits, context misses, and correctly rejected new objects are shown separately for younger and older adults in Figure 4. Follow-up pairwise analyses were conducted separately for each age group within the frontal ROIs, where the significant Condition x Age interaction effect was observed, and across age groups combined for the main effect of Condition in the right posterior ROI. There were no significant Condition x Age interaction effect (*p* = .10) or main effect of Condition (*p* = .07) observed in the left posterior ROI.

**Table 5.**
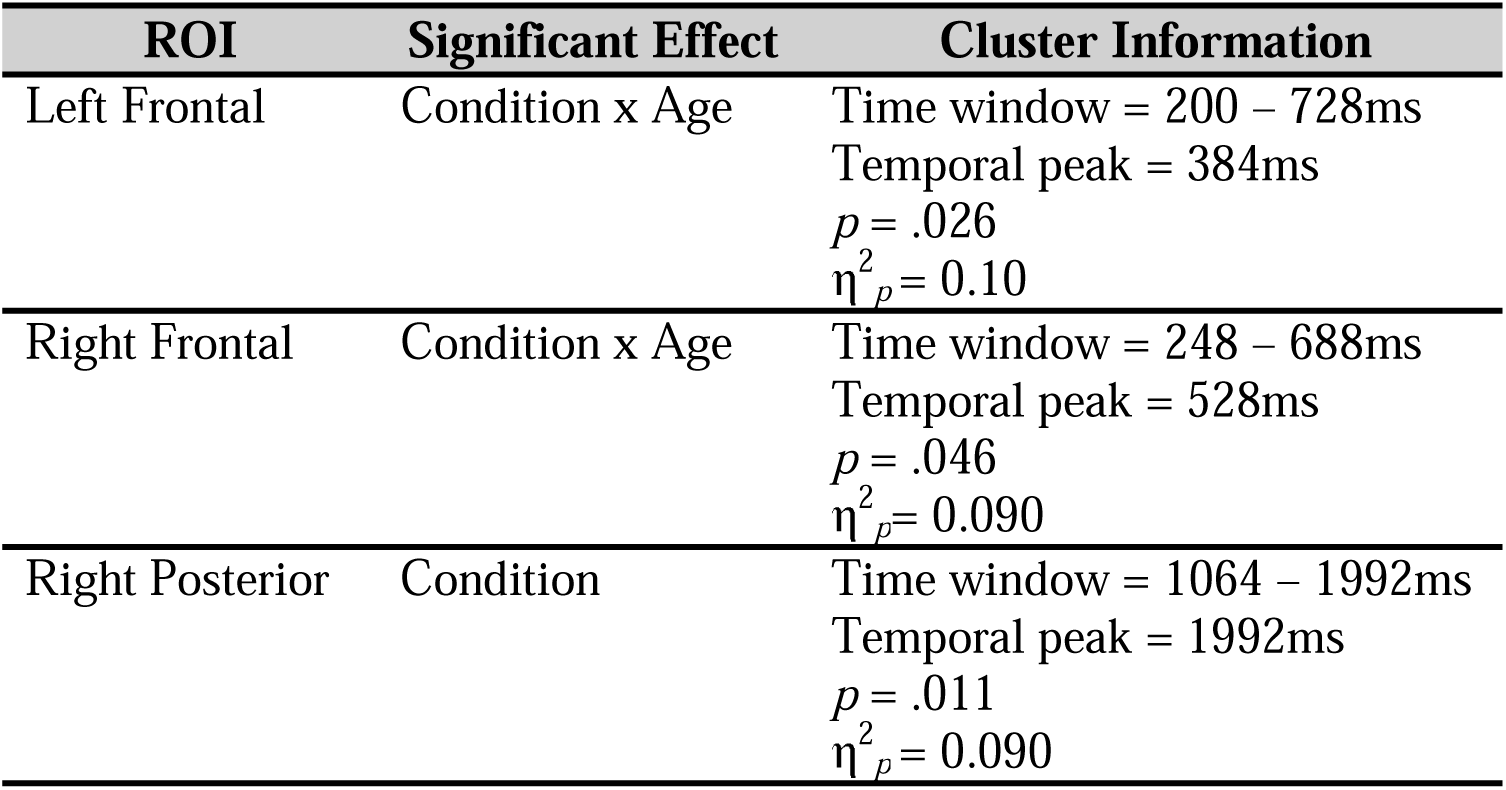
Summary of mass univariate analysis for significant clusters for matching context pairs.

| ROI | Significant Effect | Cluster Information |
| --- | --- | --- |
| Left Frontal | Condition x Age | Time window = 200 – 728ms<br>Temporal peak = 384ms<br>$p = .026$<br>$\eta^2_p = 0.10$ |
| Right Frontal | Condition x Age | Time window = 248 – 688ms<br>Temporal peak = 528ms<br>$p = .046$<br>$\eta^2_p = 0.090$ |
| Right Posterior | Condition | Time window = 1064 – 1992ms<br>Temporal peak = 1992ms<br>$p = .011$<br>$\eta^2_p = 0.090$ |

**Fig 4.**
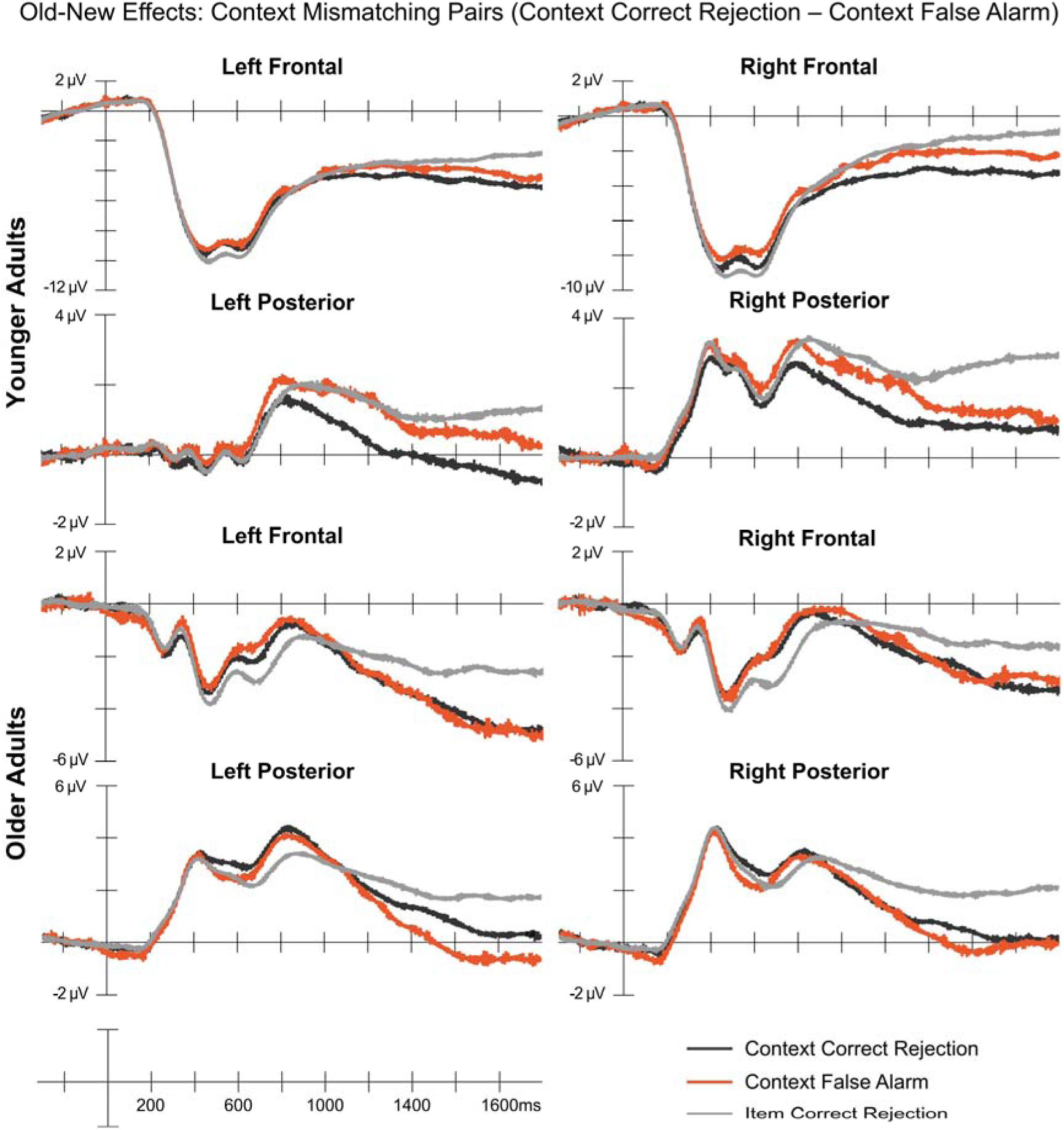
Grand ERPs for context mismatching pairs are depicted in this figure for each age group. Mismatching context pairs included average number of 59 (range: 27–98) Context Correct Rejections and 36 (range: 5–77) Context False Alarms.

Follow-up pairwise analyses in the frontal ROI for younger adults revealed early and late effects consistent with the FN400 and late frontal old-new effects, characterized by more positive-going activity for context hits and misses relative to correctly rejected new items. In the left frontal ROI, we observed significant differences between context hits and item correct rejection from 440 – 728ms with the temporal peak at 712ms (*p* = .011, η^2^*_p_* = 0.19) and between context miss and item correct rejection from 368 – 728ms with the temporal peak at 576ms (*p* = .002, η^2^*_p_* = 0.25), indicating that old objects that are correctly remembered regardless of remembering the context elicited greater positive activity than new items. There was no significant difference between context hit and context miss in this ROI (*p* = .11, η^2^*_p_*= 0.12). In the right frontal ROI, we observed significant differences between context miss and item correct rejection from 392 – 688ms with the temporal peak at 528ms (*p* = .011, η^2^*_p_* = 0.22). There were no significant differences between context hit and context miss (*p* = .096, η^2^*_p_* = 0.13) or context hit and item correct rejection (*p* = .06, η^2^*_p_* = 0.15).

The follow-up analyses for older adults in the frontal ROIs also revealed both early and late old-new effects. In the left frontal ROI, we observed a significant difference between context hit and context miss between 200 – 728ms with the temporal peak 664ms (*p* = .002, η^2^*_p_* = 0.20), between context hits and item correct rejection from 424 – 728ms with the temporal peak at 728ms (*p* = .002, η^2^*_p_*= 0.23), and between context miss and item correct rejection from 200 – 408ms with the temporal peak at 328ms (*p* = .004, η^2^*_p_* = 0.25). In the right frontal ROI, the analyses revealed significant differences between context hits and context miss from 352 – 688ms with the temporal peak at 520ms (*p* = .011, η^2^*_p_*= 0.20), between context hit and item correct rejection from 448 – 688ms with the temporal peak at 688ms (*p* = .005, η^2^*_p_* = 0.24), and between context miss and item correct rejection from 248 – 416ms with the temporal peak at 352ms (*p* = .022, η^2^*_p_* = 0.17). Notably, compared to younger adults, in addition to FN400-like effects that differentiated old and new trials, older adults showed significant differences between context hits and misses in both frontal ROIs in later time periods, suggesting that their frontal old-new effects reflected not only item familiarity but also sensitivity to whether contextual information was successfully retrieved.

The follow-up analyses for the main effect of Condition in the right posterior ROI revealed an LPN effect, with more negative-going activity for both context hit and miss relative to item correct rejection. We observed a significant difference between context hit and item correct rejection from 1320 – 1992ms with the temporal peak at 1624ms (*p* = .011, η^2^*_p_* = 0.12) and between context miss and item correct rejection from 1064 – 1992ms with the temporal peak at 1992ms (*p* = .002, η^2^*_p_* = 0.13). There were no significant differences between context hit and context miss (*p* = .16, η^2^*_p_* = 0.055) in the right posterior ROI.

Together, our findings revealed an FN400 effect in the frontal ROIs for both age groups, reflecting familiarity-based recognition for old objects relative to new items, and an LPN effect in the right posterior ROI. Notably, younger adults showed similar FN400 effects for both context hits and misses relative to new items, older adults additionally differentiated between context hits and misses in late frontal old-new effects.

##### Mismatching context pairs

Results from the mass univariate analysis for mismatching context pairs were reported in Table 6. The significant main effects indicate that old-new memory effects were present similarly across both age groups (no group interactions). Follow-up pairwise analyses were therefore conducted for significant ROIs across age groups.

**Table 6.** Summary of mass univariate analysis for significant clusters for mismatching context pairs.

| ROI | Significant Effect | Cluster Information |
| --- | --- | --- |
| Left Frontal | Condition | Time window: 1440 – 1992ms<br>Temporal peak = 1992ms<br>$p = .008$<br>$\eta^2_p = 0.11$ |
| Right Frontal | Condition | **Cluster 1:**<br>Time window = 440 – 816ms<br>Temporal peak = 720ms<br>$p = .028$<br>$\eta^2_p = 0.10$<br>**Cluster 2:**<br>Time window = 1248 – 1992ms<br>Temporal peak = 1592ms<br>$p = .012$<br>$\eta^2_p = 0.082$ |
| Left Posterior | Condition | Time window = 1144 – 1992ms<br>Temporal peak = 1992ms<br>$p = .008$<br>$\eta^2_p = 0.11$ |
| Right Posterior | Condition | Time window = 992 – 1992ms<br>Temporal peak = 1952ms<br>$p = .001$<br>$\eta^2_p = 0.13$ |

In the left frontal ROI, we observed significant differences between context correct rejection and item correct rejection from 1480 – 1992ms with the temporal peak of 1752 ms (*p* = .004 η^2^*_p_* = 0.13) and between context false alarm and item correct rejection from 1440 – 1992ms with the temporal peak at 1984ms (*p* = .001, η^2^*_p_* = 0.20). No clusters were identified in the follow-up pairwise analysis between context correct rejection and context false alarm, indicating no significant differences in this ROI.

In the right frontal ROI, we observed significant differences between context correct rejection and item correct rejection from 440 – 816ms with the temporal peak at 760ms (*p* = .001, η^2^*_p_* = 0.22) and between context false alarm and item correct rejection from 448 – 800 ms with the temporal peak at 720 ms (*p* = .009, η^2^p = 0.10). No clusters were identified between context correct rejection and context false alarm, indicating no significant differences in this ROI. A late cluster showed significant differences between context correct rejection and item correct rejection from 1512 – 1992 ms with the temporal peak at 1888 ms (*p* = .011, η^2^p = 0.11) and between context false alarm and item correct rejection from 1248 – 1992 ms with the temporal peak at 1592 ms (*p* = .001, η^2^p = 0.16). No clusters were identified in the follow-up pairwise analysis between context correct rejection and context false alarm, indicating no significant differences in this ROI.

The posterior ROIs show large LPN-like effects across age groups. In the left posterior ROI, we observed significant differences between context correct rejection and item correct rejection from 1656 – 1992 ms with the temporal peak at 1992 ms (*p* = .047, η^2^p = 0.068), between context false alarm and item correct rejection from 1144 – 1992 ms with the temporal peak at 1992 ms (*p* = .001, η^2^p = 0.263), and between context correct rejection and context false alarm from 1472 – 1960 ms with the temporal peak at 1576 ms (*p* = .022, η^2^p = 0.067). In the right posterior ROI, we observed significant differences between context correct rejection and item correct rejection from 1328 – 1992 ms with the temporal peak at 1904 ms (*p* = .002, η^2^p = 0.157) and between context false alarm and item correct rejection from 992 – 1992 ms with the temporal peak at 1984 ms (*p* = .001, η^2^p = 0.255). No clusters were identified in the follow-up pairwise analysis between context correct rejection and context false alarm, indicating no significant differences in this ROI.

In sum, our findings revealed late frontal and posterior LPN effects for mismatching context pairs across both age groups, with the LPN differentiating context correct rejections from context false alarms, consistent with recollection-based context memory retrieval, across age. To further investigate age-related differences in context memory, we conducted hierarchical regression analyses to examine whether individual differences in sleep quality and age predicted ERP memory effects.

#### Post-encoding sleep continuity is associated with matching-pair memory success effects in younger and older adults

##### Matching context pairs

Results of hierarchical regression models for Age and Sleep Continuity predicting matching pair memory effects (context hit – context miss) for ROIs and time windows with significant effects are shown in Table 5. As can be seen in Figure 5, the main effects of age reflect the different relationships between context hits and misses across age groups, especially in later time windows (600 ms and later), with more negative-going activity for context hits than misses for younger adults, and more positive-going activity for context hits than misses for older adults. Age significantly moderated the relationship between sleep continuity and these late-onsetting ERP effects for left and right frontal and left posterior ROIs. As seen in Figure 5 for the left posterior and right frontal ROIs, greater sleep continuity was associated with stronger negative-going ERP effects for younger adults (context miss > context hits) and more positive-going ERP effects for older adults (context hits > context misses).

**Table 5.** Summary of hierarchical multiple regression with age and post-encoding sleep variable predicting matching pair memory success effect.

| Time (ms) | Predictor | Block 1 |  |  | Block 2 |  |  |
| --- | --- | --- | --- | --- | --- | --- | --- |
| | | b | SE | $\beta$ | b | SE | $\beta$ |
| Left Frontal |  |  |  |  |  |  |  |
| 300 – 1100 | Age | 0.03 | 0.01 | 0.32* | 0.03 | 0.01 | 0.01* |
| 900 – 1500 | Sleep Continuity | -0.50 | 0.69 | -0.10 | -3.64 | 1.56 | -0.66* |
| 900 – 1500 | Age x Sleep Continuity | - | - | - | 0.06 | 0.03 | 0.01* |
| Right Frontal |  |  |  |  |  |  |  |
| 500 – 1000 | Age | 0.03 | 0.01 | 0.28* | 0.03 | 0.01 | 0.01* |
| 900 – 1500 | Sleep Continuity | -0.31 | 0.72 | -0.06 | -3.45 | 1.46 | -0.66* |
| 900 – 1600 | Age x Sleep Continuity | - | - | - | 0.06 | 0.03 | 0.01* |
| Left Posterior |  |  |  |  |  |  |  |
| 400 – 700 | Age | 0.02 | 0.01 | 0.32* | 0.02 | 0.01 | 0.01* |
| 800 – 1200 | Sleep Continuity | -0.16 | 0.48 | -0.04 | -2.41 | 0.98 | -0.69* |
| 800 – 1400 | Age x Sleep Continuity | - | - | - | 0.05 | 0.02 | 0.01* |

**Fig. 5.**
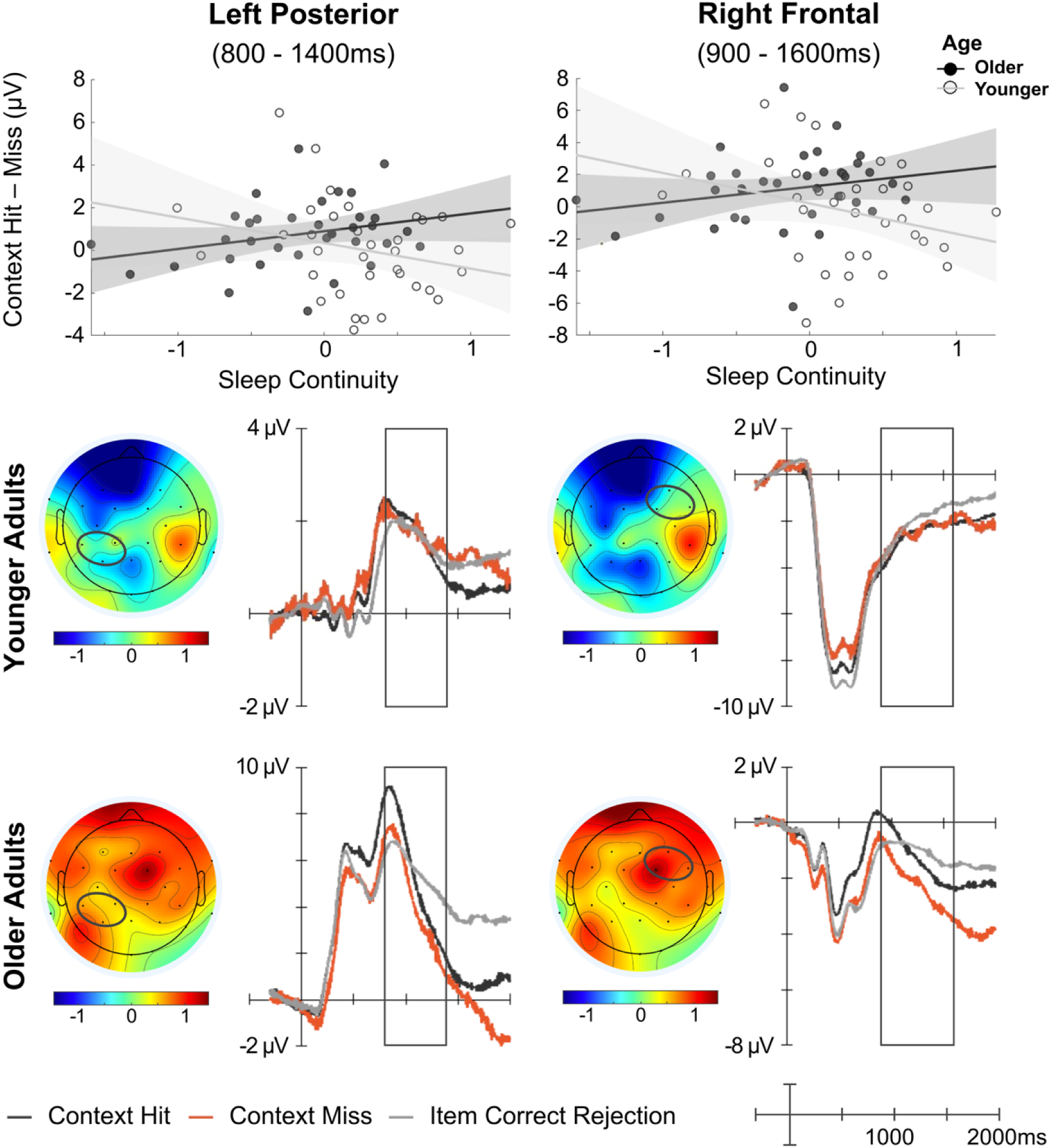
Matching pairs (Context Hit – Miss). Scatter plot of post-encoding sleep continuity against the ERP difference (μV) in the left posterior and right frontal ROIs. Scalp topographies for the highlighted windows and grand average ERPs are shown for younger and older adults.

##### Mismatching context pairs

Results for hierarchical regression models for Age and Sleep Continuity predicting Mismatching pair memory effects (Correct Rejection – False Alarm) for ROIs and time windows with significant effects are shown in Table 6. The main effect of Age was observed over posterior regions between 1700ms to 2000ms on the left and 600ms to 1200ms on the right. As can be seen in Supplemental Figure 1, the main effect of age reflects the generally more negative-going activity for context correct rejections than false alarms for young adults, and more positive-going activity for context correct rejections than false alarms for older adults. No significant relationships between sleep variables, nor interactions between sleep and age, were observed for these mismatching pair memory success ERP effects.

**Table 6.** Summary of hierarchical multiple regression with age and post-encoding sleep variables predicting Mismatching Pair Memory Success Effect.

| Time (ms) | Predictor | Block 1 |  |  | Block 2 |  |  |
| --- | --- | --- | --- | --- | --- | --- | --- |
| | | b | SE | $\beta$ | b | SE | $\beta$ |
| Left Frontal |  |  |  |  |  |  |  |
| 1700 – 2000 | Age | 0.04 | 0.02 | 0.30* | 0.04 | 0.02 | 0.01* |
| 1700 – 2000 | Sleep Continuity | 0.11 | 0.86 | 0.02 | -0.90 | 1.85 | -0.14 |
| 1700 – 2000 | Age x Sleep Continuity | - | - | - | 0.02 | 0.03 | 0.003 |
| Right Frontal |  |  |  |  |  |  |  |
| 600 – 1200 | Age | 0.02 | 0.01 | 0.31* | 0.02 | 0.01 | 0.01* |
| 600 – 1200 | Sleep Continuity | 0.45 | 0.37 | 0.16 | 0.60 | 0.81 | 0.22 |
| 600 – 1200 | Age x Sleep Continuity | - | - | - | -0.003 | 0.01 | -0.001 |

## Discussion

In the present study, we investigated whether naturalistic, post-encoding sleep quality is associated with context memory accuracy and supporting neural activity during subsequent retrieval in younger and older adults. Consistent with the existing literature, we found that younger adults showed better overall context memory performance, but older adults demonstrated greater relative retention of mismatching pairs. Post-encoding sleep continuity, which was reduced with age, was related to better context memory accuracy for mismatching pairs across both age groups. Greater post-encoding sleep continuity was associated with larger ERP differences between context hits and misses for context-matching pairs in late time periods consistent with post-retrieval monitoring and episodic reconstruction related ERPs, across age. These results and their implications are discussed below.

### Younger adults exhibited better context memory performance across delay and pair type, but older adults showed greater retention of mismatching pairs

As commonly observed in the aging literature (Cohn et al., 2008; for a review see Old & Naveh-Benjamin, 2008; Healy et al., 2005; Trelle et al., 2019; Duarte & Dulas, 2020; Kamp et al., 2022; Hokett et al., 2022; Seraji et al., 2025a; Seraji et al., 2025b), younger adults showed greater context memory accuracy than older adults for both immediate and delayed tests across matching and mismatching pair types. Across both age groups, context-matching pairs were recognized more accurately than mismatching pairs, consistent with prior studies showing greater memory for associations presented in a congruent manner between encoding and retrieval (Overman, A. A. et al. 2018). More specifically, superimposing the object on the scene context and asking participants to encode the association between the two likely facilitated their integration. By contrast, this integration would be expected to reduce recognition of mismatching object-scene pairs due to the reduced contribution of familiarity-based recognition and the need to recall the actual encoded scene for the object in order to reject the mismatching context as new (i.e. recall-to-reject) (Jones & Jacoby, 2001; Kelley & Wixted, 2001; Rotello & Heit, 2000; Rotello, Macmillan, & Van Tassel, 2004; Yonelinas, 1997, 2002).

Somewhat counterintuitively, older adults retained more mismatching pairs (i.e. forgot proportionally less) than did younger adults, with no group differences in retention of matching pairs. Prior evidence is somewhat mixed with regard to age-related differences in episodic memory retention over multi-hour to multi-day intervals (Manoli et al., 2018, Rivera-Lares et al., 2023, Studer et al., 2024). Some prior studies have shown, similar to the current results, no age-related accelerated forgetting despite impaired episodic learning as indicated by reduced immediate memory accuracy (Rivera-Lares et al., 2023, Studer et al., 2024). One recent study showed no accelerated episodic forgetting in a large sample of high-functioning older adults with reduced self-reported sleep continuity compared to that of younger adults (Studer et al., 2024), similar to the older adults in the current study who had reduced objectively measured sleep continuity compared to younger adults. One possible explanation for the preserved and somewhat elevated level of retention for older adults is that older adults’ context memory accuracy benefits more from quality sleep than does that of younger adults. That is, when of sufficient quality to support mechanisms of consolidation, post-encoding sleep may have fostered greater stability of context memory representations over time in older than younger adults. Indeed, our behavioral and ERP results show evidence of this, as discussed below.

### Greater post-encoding sleep continuity is associated with better context accuracy for mismatching pairs, across age

Greater sleep continuity during the post-encoding period was associated with better delayed memory performance for mismatching context memory pairs similarly across age groups, as we (Hokett et al., 2022; Hokett 2019) and others (Mary et al., 2013) have found in prior studies of associative memory and habitual sleep quality, though not measured post-encoding. By contrast, total sleep time was not associated with context memory retrieval. This finding aligns with prior work suggesting that quality sleep, more than sleep time, may help buffer against age-related declines in episodic memory (Sümer, E. et al 2025; Wilckens, K. A., et. al. 2014; Qin, S., et al 2023; for a review see Scullin, 2017). No significant relationship between the sleep variables and context memory performance for matching pairs was found. As we noted above, successful retrieval of mismatching object-context pairs may rely on strategic monitoring operations to a greater extent than that of matching pairs, which may be recognized on the basis of familiarity. Specifically, the ability to recall prior episodic associations in order to reject mismatching new ones necessitates PFC-dependent executive control functions that are impaired in aging, including post-retrieval monitoring and episodic reconstruction that operate on recollected memory contents (Duarte, A., & Dulasa, M. R. 2020). As familiarity-based recognition is less hippocampal-dependent than is recollection, it may benefit less from hippocampal-dependent consolidation mechanisms engaged during sleep (Rasch & Born, 2013). As a result, sleep quality may show weaker associations with context matching than context mismatching pairs across age groups.

### Sleep continuity is associated with matching pair memory success ERP effects, across age

Individuals with greater post-encoding sleep continuity showed larger ERP differences between context hits and misses for context-matching pairs in late time periods (800-1600 ms) over frontal and posterior scalp sites. The time course of these effects and their sensitivity to episodic memory accuracy are consistent with late frontal and LPN ERPs, implicated in post-retrieval monitoring (Cruse and Wilding, 2009; Dulas and Duarte, 2013; Wilding and Rugg, 1996) and episodic reconstruction, respectively (for a review see: Mecklinger et al., 2016). Interestingly, these associations were moderated by age, due to a polarity difference between memory conditions, with more negative-going activity for context hits than misses in younger adults and more positive-going activity for context hits than misses in older adults (see Fig.5). While this age effect was not predicted, it is somewhat similar to a pattern we previously observed in an object-scene context memory retrieval task in which late negative-going ERPs were pronounced, potentially obscuring the typically positive-going late frontal old-new effects (James et al. 2016).

Given these post-retrieval mechanisms are likely to be particularly engaged during retrieval of mismatching pairs, as discussed earlier, it was somewhat surprising that sleep continuity was not associated with these late ERPs for mismatching pairs. While it can be difficult to interpret null results, especially given the positive associations with mismatching context memory accuracy, we see two related possible explanations. First, robust engagement of these ERPs for mismatching, relative to matching trials, is consistent with higher demands on these processes across age groups. This relatively maximal engagement across participants may have obscured the detection of individual differences in these operations and/or they were recruited by the demands of the task regardless of sleep quality. It is also possible that the positive associations between post-encoding sleep quality and mismatching pair memory accuracy reflect consolidation mechanisms operating during this post-encoding period, including hippocampal-dependent reactivation of encoded representations during NREM, which actigraphy isn’t sensitive to. Future studies could include EEG and fMRI monitoring during post-encoding sleep would be needed to assess this possibility.

## Conclusion

In conclusion, our findings suggest that better naturalistic, post-encoding sleep quality promotes neural processes underlying memory of item-context associations and the successful post-retrieval discrimination of context hits from context misses, across young and older adults. These findings are consistent with the broader literature on sleep-dependent memory consolidation, suggesting that more continuous sleep may facilitate the consolidation of associative memory that are especially vulnerable to age-related decline. Importantly, these results highlight post-encoding sleep quality as a potentially modifiable factor that may help attenuate age-related episodic memory deficits. A notable strength of the present study is the use of actigraphy-based sleep measurement over multiple nights in at-home sleep environment and measurement of context memory retrieval-related neural activity after a period of sleep at home, which together provide a more ecologically valid characterization of the sleep-memory relationship than prior laboratory-based sleep manipulation studies. Future research should examine these relationships across a broader lifespan sample with larger and more diverse participant groups to better characterize how sleep continuity influences memory consolidation across different stages of aging.

## Data availability statement

The data from the current study are publicly available on OSF (https://osf.io/t8vk3/overview?view_only=4e4d3f0033e249a8ad832354544897cf)

## Author contributions

Chuu Nyan: Data curation, Formal analysis, Investigation, Project administration, Visualization, Writing – original draft preparation, and Writing – review & editing

AJ Wachnin: Data curation, Formal analysis, Investigation, Project administration, Visualization, Writing – original draft preparation, and Writing – review & editing

Soroush Mirjalili: Formal analysis, Writing – review & editing

Sahana Ram: Investigation, Writing – review & editing

Masoud Seraji: Methodology, Software, Writing – review & editing

Audrey Duarte: Conceptualization, Funding acquisition, Methodology, Resources, Supervision, Writing – Original Draft Preparation, and Writing – Review & Editing

## Acknowledgements

We thank all our research participants for their participation.

## Diversity in Citation Practices

The gender balance of this article’s reference list was estimated using the GCBI-alyzer tool (Fulvio et al., 2021), which predicts the gender of each reference’s first and last author from first names. The categorizable references are 43.6% man/man, 25.5% woman/man, 12.7% man/woman, and 18.2% woman/woman (first author/last author). Relative to Journal of Cognitive Neuroscience base rates, the corresponding gender citation balance indices are +0.072, −0.171, +0.097, and +0.063, respectively (positive indicates over-citation, negative indicates under-citation).

---

**Supplemental Fig. 1.**
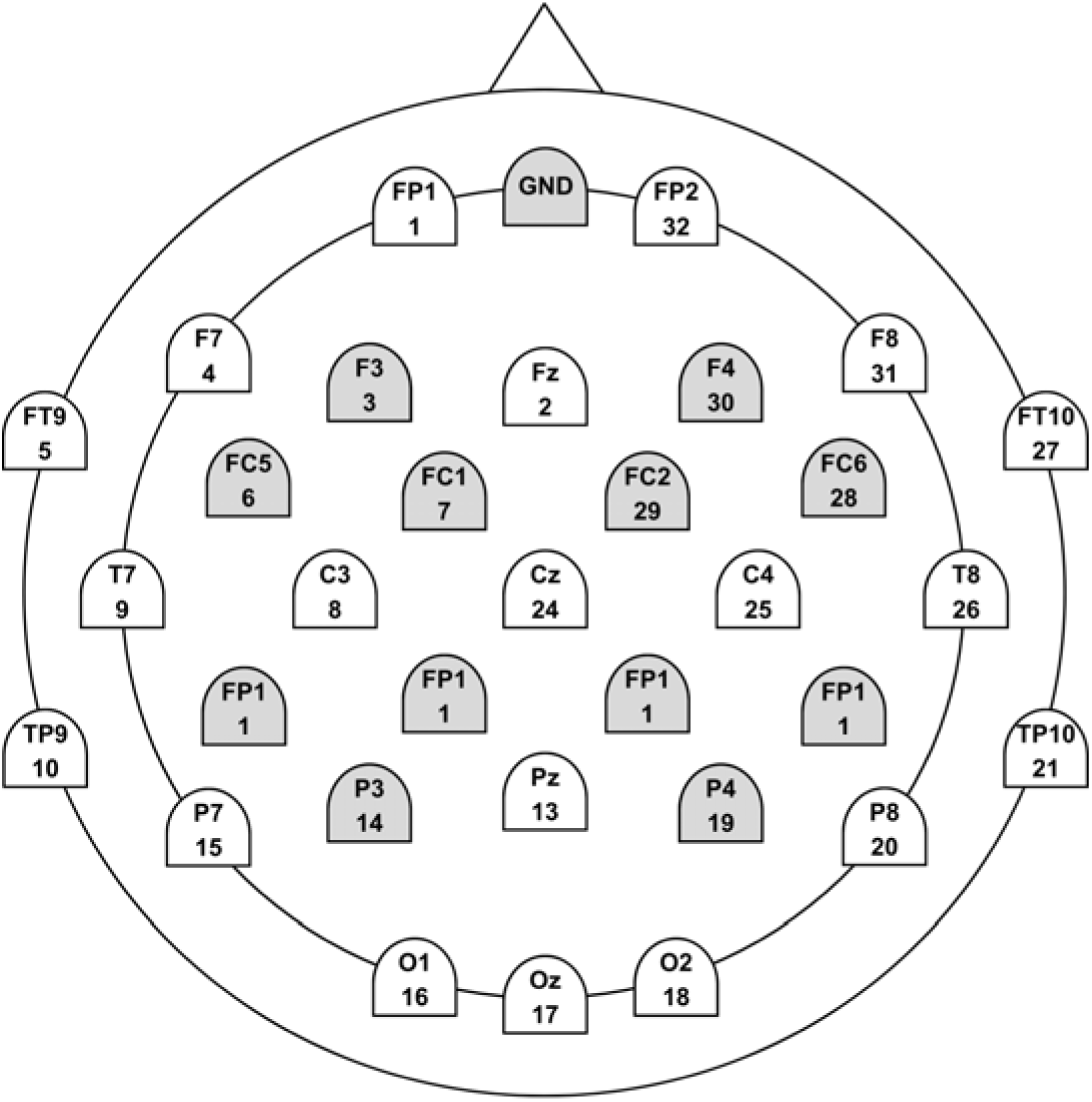
ERP regions of interest (ROIs): Left Frontal (F3, FC1, FC5), Right Frontal (F4, FC2, FC6), Left Posterior (CP1, CP5, P3), Right Posterior (CP2, CP6, P4).

