## Supplemental for "Individual differences in post-encoding sleep continuity predict context memory accuracy and supporting ERPs in younger and older adults"

**Supplemental Findings**

Supplemental Table 1. Mean sleep variables resulted in two sleep components: Sleep Continuity (Sleep Efficiency, Number of Awakenings, WASO, Sleep Fragmentation Index) and Sleep Time (TST).

| **Sleep variables** | | **Whole sleep period**  **(168 hours)** | | **Pre-encoding**  **(72 hours)** | |
| --- | --- | --- | --- | --- | --- |
|  |  | **Sleep Continuity** | **Sleep Time** | **Sleep Continuity** | **Sleep Time** |
| Mean | Total Sleep Time | -0.03 | 0.97 | -0.03 | 0.97 |
|  | Sleep Efficiency | -0.84 | 0.29 | -0.84 | 0.29 |
|  | Number of Awakening | 0.82 | 0.41 | 0.82 | 0.41 |
|  | Wake After Sleep Onset (WASO) | 0.88 | 0.32 | 0.88 | 0.32 |
|  | Sleep Fragmentation Index | 0.86 | -0.26 | 0.86 | -0.26 |

Supplemental Table 2. Summary of hierarchical multiple regression with age and pre-encoding sleep predicting delayed context memory accuracy.

| **Outcome variable: Memory Retention Matching Pairs** | | | | | | |
| --- | --- | --- | --- | --- | --- | --- |
| Predictor variables | **Block 1** | | | **Block 2** | | |
|  | b | SE | *β* | b | SE | *β* |
| Intercept | -0.09 | 0.02 |  | -0.08 | 0.02 |  |
| Age | 0.04 | 0.03 | 0.21 | 0.04 | 0.03 | 0.21 |
| Sleep continuity | -0.01 | 0.03 | -0.03 | -0.003 | 0.03 | -0.02 |
| Age x Sleep Continuity |  |  |  | -0.01 | 0.06 | -0.03 |
| R^2^ | 0.04 | | | 0.04 | | |
| ∆R^2^ | 0.04 | | | <0.001 | | |
| F for ∆R^2^ | 1.38 | | | 0.92 | | |
| **Outcome variable: Memory Retention Matching Pairs** | | | | | | |
| Intercept | -0.09 | 0.02 |  | -0.09 | 0.02 |  |
| Age | 0.04 | 0.02 | 0.20 | 0.04 | 0.02 | 0.20 |
| Sleep time | 0.02 | 0.01 | 0.16 | 0.01 | 0.01 | 0.08 |
| Age x Sleep Time |  |  |  | 0.02 | 0.02 | 0.13 |
| R^2^ | 0.07 | | |  | 0.08 |  |
| ∆R^2^ | 0.07 | | |  | 0.01 |  |
| F for ∆R^2^ | 2.35 | | |  | 1.80 |  |
| **Outcome variable: Memory Retention Mismatching Pairs** | | | | | | |
| Intercept | -0.05 | 0.02 |  | -0.04 | 0.02 |  |
| Age | -0.11 | 0.03 | -0.40* | -0.11 | 0.03 | -0.40* |
| Sleep Continuity | <0.001 | 0.03 | -0.001 | -0.003 | 0.04 | -0.01 |
| Age x Sleep Continuity |  |  |  | 0.01 | 0.07 | 0.01 |
| R^2^ | 0.16 | | | 0.16 | | |
| ∆R^2^ | 0.16 | | | <0.001 | | |
| F for ∆R^2^ | 6.26 | | | 4.12 | | |
| **Outcome variable: Memory Retention Mismatching Pairs** | | | | | | |
| Intercept | -0.05 | 0.02 |  | -0.05 | 0.02 |  |
| Age | -0.11 | 0.03 | -0.40* | -0.11 | 0.03 | -0.40* |
| Sleep Time | -0.003 | 0.02 | -0.02 | -0.003 | 0.02 | -0.02 |
| Age x Sleep Time |  |  |  | 0.002 | 0.03 | 0.01 |
| R^2^ | 0.16 | | | 0.16 | | |
| ∆R^2^ | 0.16 | | | <0.001 | | |
| F for ∆R^2^ | 6.28 | | | 4.13 | | |

Supplemental Table 3. Summary of hierarchical multiple regression with age and post-encoding sleep predicting context memory retention.

| **Outcome variable: Memory Retention Matching Pairs** | | | | | | |
| --- | --- | --- | --- | --- | --- | --- |
| Predictor variables | **Block 1** | | | **Block 2** | | |
|  | b | SE | *β* | b | SE | *β* |
| Intercept | -0.09 | 0.02 |  | -0.09 | 0.02 |  |
| Age | 0.04 | 0.03 | 0.22 | 0.04 | 0.03 | 0.22 |
| Sleep continuity | -0.01 | 0.02 | -0.06 | -0.01 | 0.03 | -0.05 |
| Age x Sleep Continuity |  |  |  | -0.004 | 0.05 | -0.02 |
| R^2^ | 0.04 | | | 0.04 | | |
| ∆R^2^ | 0.04 | | | <0.001 | | |
| F for ∆R^2^ | 1.47 | | | 0.97 | | |
| **Outcome variable: Memory Retention Matching Pairs** | | | | | | |
| Intercept | -0.09 | 0.02 |  | -0.09 | 0.02 |  |
| Age | 0.04 | 0.02 | 0.21 | 0.04 | 0.02 | 0.21 |
| Sleep time | 0.02 | 0.01 | 0.16 | 0.01 | 0.02 | 0.14 |
| Age x Sleep Time |  |  |  | 0.005 | 0.2 | 0.04 |
| R^2^ | 0.06 | | | 0.07 | | |
| ∆R^2^ | 0.06 | | | -0.001 | | |
| F for ∆R^2^ | 2.34 | | | 1.55 | | |
| **Outcome variable: Memory Retention Mismatching Pairs** | | | | | | |
| Intercept | -0.04 | 0.02 |  | -0.04 | 0.02 |  |
| Age | -0.11 | 0.03 | -0.41* | -0.11 | 0.04 | -0.39* |
| Sleep Continuity | 0.01 | 0.03 | 0.02 | 0.03 | 0.04 | 0.10 |
| Age x Sleep Continuity |  |  |  | -0.05 | 0.07 | -0.11 |
| R^2^ | 0.16 | | | 0.16 | | |
| ∆R^2^ | 0.16 | | | -0.01 | | |
| F for ∆R^2^ | 6.28 | | | 4.32 | | |
| **Outcome variable: Memory Retention Mismatching Pairs** | | | | | | |
| Intercept | -0.05 | 0.02 |  | -0.05 | 0.02 |  |
| Age | -0.11 | 0.03 | -0.40* | -0.11 | 0.03 | -0.40* |
| Sleep Time | 0.002 | 0.02 | 0.01 | 0.001 | 0.02 | 0.01 |
| Age x Sleep Time |  |  |  | 0.001 | 0.03 | 0.004 |
| R^2^ | 0.16 | | | 0.16 | | |
| ∆R^2^ | 0.16 | | | <0.001 | | |
| F for ∆R^2^ | 6.27 | | | 4.12 | | |

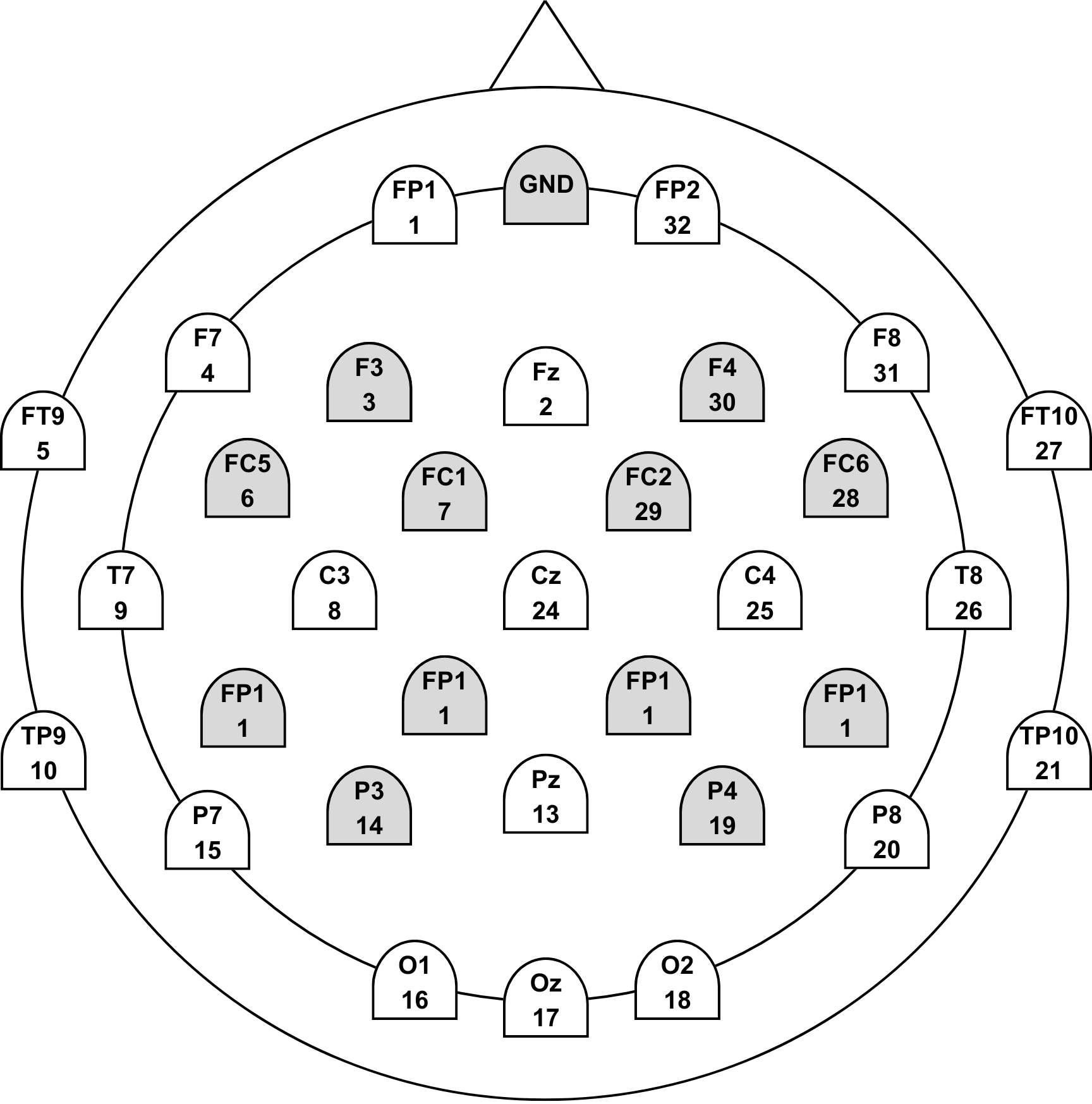

**Supplemental Fig. 1.** ERP regions of interest (ROIs): Left Frontal (F3, FC1, FC5), Right Frontal (F4, FC2, FC6), Left Posterior (CP1, CP5, P3), Right Posterior (CP2, CP6, P4).
